# Ultra-fast genotyping of SNPs and short indels using GPU acceleration

**DOI:** 10.1101/2023.04.14.534526

**Authors:** Jørgen Wictor Henriksen, Knut Dagestad Rand, Geir Kjetil Sandve, Ivar Grytten

**Author notes:** Correspondence to Ivar Grytten < >.

## Abstract

As decreasing DNA sequencing costs leads to a steadily increasing rate of generated data, the development of efficient algorithms for processing of the sequence data is increasingly important to reduce costs and energy consumption. Recent work have shown that genotyping can be done efficiently and accurately using alignment-free methods that are based on analyzing kmers from sequenced reads. In particular, we have recently presented the KAGE genotyper, which uses an efficient pangenome representation of known individuals in a population to further increase accuracy and efficiency. While existing genotypers like KAGE use the Central Processing Unit (CPU) to count and analyze kmers, the Graphical Processing Unit (GPU) has shown promising for reducing runtime for similar problems.

We here present GKAGE, a new and improved version of KAGE that utilizes the GPU to further increase the computational efficiency. This is done by counting and analyzing large amounts of kmers in the many parallel cores of a GPU. We show that GKAGE is, on hardware of comparable cost, able to genotype an individual up to an order of magnitude faster than KAGE while producing the same output, which makes it by far the fastest genotyper available today. GKAGE can run on consumer-grade GPUs, and enables genotyping of a human sample in only a matter of minutes without the need for expensive high-performance computers. GKAGE is open source and available at https://github.com/kage-genotyper/kage.

## Introduction

The cost of sequencing a full human genome has fallen drastically in recent years, and consumers can now get their whole genome sequenced for a few hundred dollars [1], a fraction of what was the price only a few years ago. As millions of genomes are likely to be sequenced in the coming years, there is an ever-increasing need for efficient methods for analyzing this genomic data. At the core of such analysis is variant detection, determining which genetic variants are present in a sample based on the sequenced reads.

Recent methods [2,3,4,5] have shown that detecting variants in a human sample can be performed efficiently and with high accuracy by genotyping the sample against an existing database of known human variation. Such methods use prior knowledge from e.g. the 1000 Genomes Project [6] about where in the genome individuals frequently have variation, and for each such genetic variant use the genomic reads from the sample to infer the most likely genotype. While such methods have traditionally been based on aligning reads to a reference genome, which is slow, recent alignment-free methods have shown that drastic speedup can be achieved by instead analyzing kmers from the sequenced reads. In a recent publication [2], we proposed a highly efficient alignment-free method, KAGE, that uses prior knowledge from a population to achieve high genotyping accuracy while being more computationally efficient than other alignment-free genotypers. Alignment-free genotypers, like KAGE, generally rely on analyzing kmers from reads against kmers associated with genomic variants.

Prior to genotyping, kmers that represent alleles of variants of interest are collected and stored in an index (e.g. a hashmap) that enables lookup of kmers from reads to variant alleles. This is typically done once for a set of variants, and this index can then be reused for genotyping any individual against those variants. When genotyping an individual, kmers from reads are collected and looked up in the kmer index, to obtain kmer counts for each variant allele. Generally, genotype probabilities are then found by analyzing the counts of how many reads support the various alleles of the variants of interest. In KAGE, these counts are combined with expected counts sampled from a population to obtain better estimates.

In contrast to other genotypers, especially alignment-based genotypers like GATK [7], the time-consuming steps of KAGE (like processing and counting kmers) can readily benefit from the massive parallelization that the Graphical Processing Unit (GPU) offers [9]. Here, we show that applying GPU acceleration to the compute-heavy parts of KAGE leads to a genotyper GKAGE of unsurpassed computational efficiency. GKAGE uses the GPU to process genomic reads, extract kmers and count the number of kmers that support alleles of genetic variants. We show that this results in a substantial speedup of up to 10X over the original KAGE genotyper (which was already faster than any other genotyper), while producing the exact same output. GKAGE has been implemented so that it is able to run even on standard consumer GPUs, and is able to genotype a whole human sample in a matter of minutes.

## Results

GKAGE is a GPU-accelerated version of the recently published KAGE genotyper. GKAGE uses CUDA-enabled GPUs to efficiently parse and encode kmers from reads, to genotype a set of known SNPs and indels based on the kmer counts. GKAGE produces output that is identical to that of KAGE, with reduced runtime on systems that support the CUDA interface for GPU acceleration. The software is open source and available at https://github.com/kage-genotyper/kage. As part of GKAGE, we have also implemented a static GPU hashmap for counting kmers through a Python interface, available at https://github.com/kage-genotyper/cucounter.

We have recently shown that KAGE is an order of magnitude faster than existing genotypers while giving better or comparable accuracy [2]. We thus only benchmark GKAGE against KAGE to show the effect of GPU acceleration. We do this by running GKAGE and KAGE on a human whole genome sample (15x coverage) on two different systems:

1. A high-performance server with an AMD EPYC 7742 64-Core CPU and two NVIDIA Tesla V100 GPUs. KAGE was run using 16 threads and GKAGE was run using one GPU.

2. A regular desktop computer with an 11th Gen Intel(R) Core(TM) i5-11400F @ 2.60GHz CPU with 6 cores and a NVIDIA GTX 1660 super GPU. KAGE was run using 6 threads.

Table 1 shows the runtimes on these two systems. GKAGE is approximately 5x faster on the high-end system and more than 10x faster on the desktop computer. Since GKAGE only needs the kmer counts of a predefined set of kmers (those associated with variants), and no existing GPU-based kmer counter is able to count only a given set of kmers, we have implemented our own kmer counter as part of GKAGE. An alternative solution would be to count all kmers using an existing tool and filter out those kmers that are relevant. Table 1 also shows the time spent by the GPU kmer counter Gerbil [8] to only count kmers.

**Table 1:** Running times of KAGE, GKAGE and Gerbil (only kmer counting)

|  | KAGE | GKAGE | Gerbil (only kmer counting) |
| --- | --- | --- | --- |
| Desktop computer | 1993 sec | 178 sec | 464 sec |
| High-performance computer | 510 sec | 94 sec | 438 sec |

## Methods

### Implementation

Here we describe more in detail how GKAGE has been implemented. While GKAGE is implemented as part of KAGE, and shares large parts of its code with KAGE, compute-heavy parts have been reimplemented so that the GPU is utilized. GKAGE implements GPU support for two bottleneck components of KAGE that were suitable for GPU acceleration:

#### Reading and encoding kmers

from a FASTA/FASTQ file is achieved in KAGE by using BioNumPy [10], a Python library built on top of NumPy [11]. BioNumPy uses NumPy to efficiently read chunks of DNA reads from fasta files, encode the bases to a 2-bit representation, and then encode the valid kmers as 64-bit integer representations in an array. GPU support for this step was achieved by utilizing CuPy [12], a GPU accelerated computing library with an interface that closely follows that of NumPy. This component was implemented in GKAGE by replacing the NumPy module in BioNumPy with CuPy, effectively replacing all NumPy function calls with calls to CuPy’s functions providing the same functionality with GPU acceleration. This strategy worked out of the box for most parts of the BioNumPy solution, with only a few custom modifications having to be made due to certain functions in NumPy’s interface not being supported by CuPy.

#### Counting kmers

As part of GKAGE, we have implemented a static hashtable for counting a predefined set of kmers on the GPU. The implementation supports parallel and high-throughput hashing and counting of large chunks of kmers simultaneously on the GPU. This static hashtable is implemented as a C++ class in CUDA [13], with two arrays of 64-bit and 32-bit unsigned integers to represent kmers (keys) and counts (values), respectively. CUDA kernels are implemented that handle insertion (only once during initialization of the hashtable), counting and querying of kmers. The hashtable uses open addressing and a simple linear probing scheme with a murmur hash [14] for the keys.

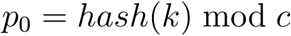

To find the position of a kmer *k* in the hashtable, the initial probe position *po* is found by computing

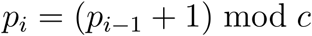

where *hash* is a murmur hash function and *c* is the capacity of the hashtable. If is occupied by a different kmer than, the next probing position can be computed given the previous probing position *p_i-1_* with

The probing will continue until either *k* or an empty slot in the hashtable is observed (See Figure 1 for an illustration of this).

**Figure 1:**
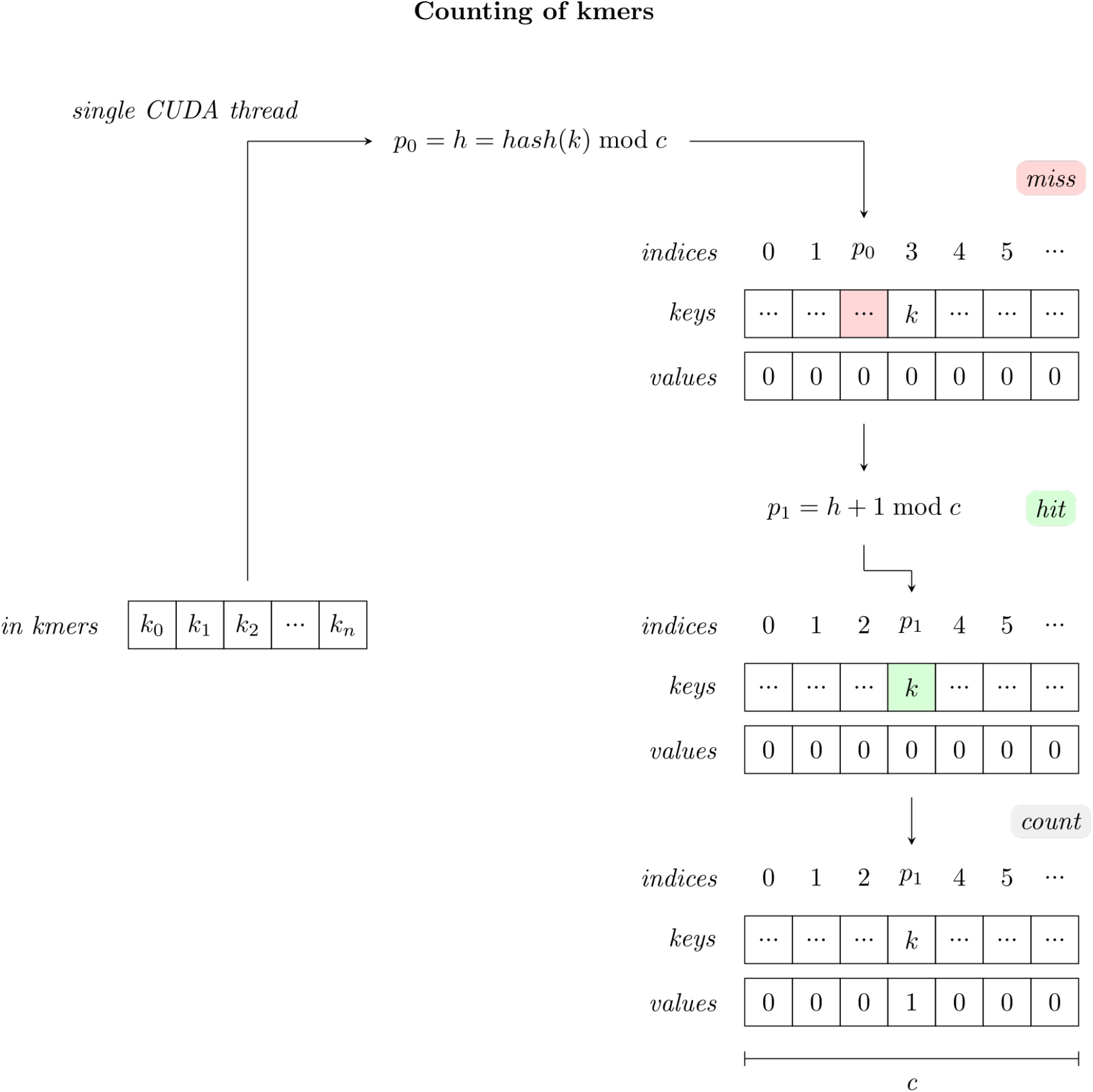
As an array of 64-bit integer encoded kmers are counted by the hash table, each CUDA thread will compute the first probe position *p_0_* for each individual kmer, and then continue probing by linearly moving up to the next consecutive slot until either an empty slot or the original kmer handled by the thread is observed. If an empty slot is observed, the thread terminates. If the original kmer is observed, the value at the current slot is increased.

The hashtable supports three main operations: insertion, counting and querying. In each of the cases, the input is an array of 2-bit encoded kmers. When querying, the return value is an array of counts associated with the input kmers. For insertion, counting or querying of *n* kmers, *n* CUDA threads are launched. Each thread is then responsible for fulfilling the relevant operation associated with the kmer, i.e. incrementing or fetching the count associated with the kmer in the hashtable, all achieved using the probing scheme previously described. Furthermore, for insertions and count updates, CUDA atomic operations are used to avoid race conditions. To use the hashtable class in Python, C++ bindings are implemented using pybind11 [15].

Since KAGE only needs to count kmers that are preselected to represent alleles of known variants, which typically is only a small subset of the kmers present in genomic reads, the hashmap needed for this requires only a few gigabytes of memory. Thus, when genotyping 28 million variants of a human sample, a GPU with 4GB of memory is sufficient (see details below).

### Memory Usage

The memory usage for GKAGE is influenced by two factors: The size of the kmer index, and the number of sequences to handle at a time (chunk size). Since the chunk size can be set to a suitable number for a given system (we used 10 MB in our experiments), the main factor will be the size of the kmer-index.

The size of the kmer-index itself is again influenced by three factors: The number of variants to be genotyped (*N_υ_*), the average number of kmers per variant (*k*), and the loading factor of the hash table (*L*). Given these, the memory will be in the order of *O*(*N_υ_* * *K*/*L*). Thus, for a fixed loading factor (default 0.57), the memory consumption of GKAGE is primarily driven by the number of variants to be genotyped. For systems with little memory, the loading factor can be increased to remove the memory consumption, but that can severely impede performance as the loading factor approaches 1. For genotyping the 28 million variants in the experiment in table 1, with an average of 3.4 kmers per variant and a loading factor of, the memory consumption of the index is (28 * 10^6^ * 3.4/0.57) * (8 + 4)*B* = 2*GB* (using 8 bytes for keys and 4 bytes for counts).

### Benchmarks

Benchmarking was performed on two different computer systems, as described in the Results section, using simulated reads for a whole genome sample with 15x coverage. A Snakemake [16] pipeline for reproducing the benchmarking results can be found at https://github.com/kage-genotyper/GKAGE-benchmarking.

## Discussion

We have presented GKAGE, a GPU accelerated, alignment-free genotyper based on a previously published CPU-based genotyper KAGE. Our results show that alignment-free genotyping is an ideal problem for GPU acceleration. While the existing KAGE genotyper is already fast by today’s standards, GKAGE is considerably faster, enabling rapid genotyping even on consumer-grade computers. We see these improvements of computational efficiency as highly beneficial considering the continually decreasing cost and expanding capacity at the experimental side of whole-genome sequencing.

Since the original KAGE genotyper was implemented mainly using the array programming libraries NumPy and BioNumPy in Python, GPU support could be added to the existing code base in a clean way by using the CuPy library combined with some custom CUDA kernels with Python wrappers. We thus see our work as a strong example of how the addition of GPU support to existing tools is typically highly feasible and beneficial in cases where many independent operations are performed on an array of data, which is common for problems in computational biology. As GPUs are becoming steadily cheaper and more available, we thus see a huge potential in improving the computational efficiency of existing methods and tools, which in many cases can be achieved quite easily through the Python ecosystem with packages such as CuPy [12], Numba [17] and BioNumPy [10].

## Notes

### Competing Interest Statement

The authors have declared no competing interest.

## References

1. A New Wave of Genomics for All Diana Crow Cell (2019-03) https://doi.org/gfw9g5 DOI: 10.1016/j.cell.2019.02.041 · PMID: 30901548

2. KAGE: fast alignment-free graph-based genotyping of SNPs and short indels Ivar Grytten, Knut Dagestad Rand, Geir Kjetil Sandve Genome Biology (2022-10-04) https://doi.org/grpzvf DOI: 10.1186/s13059-022-02771-2 · PMID: 36195962 · PMCID: PMC9531401

3. MALVA: Genotyping by Mapping-free ALlele Detection of Known VAriants Luca Denti, Marco Previtali, Giulia Bernardini, Alexander Schönhuth, Paola Bonizzoni iScience (2019-08) https://doi.org/gmhw28 DOI: 10.1016/j.isci.2019.07.011 · PMID: 31352182 · PMCID: PMC6664100

4. Pangenome-based genome inference allows efficient and accurate genotyping across a wide spectrum of variant classes Jana Ebler, Peter Ebert, Wayne E Clarke, Tobias Rausch, Peter A Audano, Torsten Houwaart, Yafei Mao, Jan O Korbel, Evan E Eichler, Michael C Zody, … Tobias Marschall Nature Genetics (2022-04) https://doi.org/grp6v6 DOI: 10.1038/s41588-022-01043-w · PMID: 35410384 · PMCID: PMC9005351

5. Graphtyper enables population-scale genotyping using pangenome graphs Hannes P Eggertsson, Hakon Jonsson, Snaedis Kristmundsdottir, Eirikur Hjartarson, Birte Kehr, Gisli Masson, Florian Zink, Kristjan E Hjorleifsson, Aslaug Jonasdottir, Adalbjorg Jonasdottir, … Bjarni V Halldorsson Nature Genetics (2017-09-25) https://doi.org/gbx7v6 DOI: 10.1038/ng.3964 · PMID: 28945251

6. A global reference for human genetic variation, Adam Auton, Gonçalo R Abecasis, David M Altshuler (Co-Chair), Richard M Durbin (Co-Chair), Gonçalo R Abecasis, David R Bentley, Aravinda Chakravarti, Andrew G Clark, Peter Donnelly, … Nature (2015-09-30) https://doi.org/73d DOI: 10.1038/nature15393 · PMID: 26432245 · PMCID: PMC4750478

7. Scaling accurate genetic variant discovery to tens of thousands of samples Ryan Poplin, Valentin Ruano-Rubio, Mark A DePristo, Tim J Fennell, Mauricio O Carneiro, Geraldine A Van der Auwera, David E Kling, Laura D Gauthier, Ami Levy-Moonshine, David Roazen, … Eric Banks Cold Spring Harbor Laboratory (2017-11-14) https://doi.org/ggmrvr DOI: 10.1101/201178

8. Gerbil: a fast and memory-efficient k-mer counter with GPU-support Marius Erbert, Steffen Rechner, Matthias Müller-Hannemann Algorithms for Molecular Biology (2017-03-31) https://doi.org/gkzhfr DOI: 10.1186/s13015-017-0097-9 · PMID: 28373894 · PMCID: PMC5374613

9. GPU Acceleration of Advanced k-mer Counting for Computational Genomics Huiren Li, Anand Ramachandran, Deming Chen 2018 IEEE 29th International Conference on Application-specific Systems, Architectures and Processors (ASAP) (2018-07) https://doi.org/grp3kq DOI: 10.1109/asap.2018.8445084

10. BioNumPy: Fast and easy analysis of biological data with Python Knut Rand, Ivar Grytten, Milena Pavlovic, Chakravarthi Kanduri, Geir Kjetil Sandve Cold Spring Harbor Laboratory (2022-12-22) https://doi.org/grp3k6 DOI: 10.1101/2022.12.21.521373

11. Array programming with NumPy Charles R Harris, KJarrod Millman, Stéfan J van der Walt, Ralf Gommers, Pauli Virtanen, David Cournapeau, Eric Wieser, Julian Taylor, Sebastian Berg, Nathaniel J Smith, … Travis E Oliphant Nature (2020-09-16) https://doi.org/ghbzf2 DOI: 10.1038/s41586-020-2649-2 · PMID: 32939066 · PMCID: PMC7759461

12. CuPy: NumPy/SciPy-compatible Array Library for GPU-accelerated Computing with Python Ryosuke Okuta, Yuya Unno, Daisuke Nishino, Shohei Hido, Crissman Loomis Thirty-first Annual Conference on Neural Information Processing Systems (NeurIPS)

13. CUDA C++ Programming Guide NVIDIA Corporation & Affiliates (2023) https://docs.nvidia.com/cuda/cuda-c-programming-guide/#

14. MurmurHash3 Austin Appleby (2023) https://github.com/aappleby/smhasher/blob/master/src/MurmurHash3

15. pybind11 -- Seamless operability between C++11 and Python Wenzel Jakob GitHub (2023) https://github.com/pybind/pybind11

16. Sustainable data analysis with Snakemake Felix Mölder, Kim Philipp Jablonski, Brice Letcher, Michael B Hall, Christopher H Tomkins-Tinch, Vanessa Sochat, Jan Forster, Soohyun Lee, Sven O Twardziok, Alexander Kanitz, … Johannes Köster F1000Research (2021-04-19) https://doi.org/gj76rq DOI: 10.12688/f1000research.29032.2 · PMID: 34035898 · PMCID: PMC8114187

17. Numba Siu Kwan Lam, Antoine Pitrou, Stanley Seibert Proceedings of the Second Workshop on the LLVM Compiler Infrastructure in HPC (2015-11-15) https://doi.org/gf3nks DOI: 10.1145/2833157.2833162

